# Tendon-derived biomimetic surface topographies induce phenotypic maintenance of tenocytes *in vitro*

**DOI:** 10.1101/2020.07.23.217224

**Authors:** Aysegul Dede Eren, Aliaksei Vasilevich, E. Deniz Eren, Phanikrishna Sudarsanam, Urandelger Tuvshindorj, Jan de Boer, Jasper Foolen

## Abstract

The tenocyte niche contains biochemical and biophysical signals that are needed for tendon homeostasis. The tenocyte phenotype is correlated with cell shape *in vivo* and *in vitro*, and shape-modifying cues are needed for tenocyte phenotypical maintenance. Indeed, cell shape changes from elongated to spread when cultured on a flat surface, and rat tenocytes lose the expression of phenotypical markers throughout five passages. We hypothesized that tendon gene expression can be preserved by culturing cells in the native tendon shape. To this end, we reproduced the tendon topographical landscape into tissue culture polystyrene, using imprinting technology. We confirmed that the imprints forced the cells into a more elongated shape, which correlated with the level of Scleraxis expression. When we cultured the tenocytes for seven days on flat surfaces and tendon imprints, we observed a decline in tenogenic marker expression on flat but not on imprints. This research demonstrates that native tendon topography is an important factor contributing to the tenocyte phenotype. Tendon imprints therefore provide a powerful platform to explore the effect of instructive cues originating from native tendon topography on guiding cell shape, phenotype and function of tendon-related cells.

## Introduction

Tendon is a unique type of connective tissue that transmits muscle contraction forces to bones to produce motion and maintain body posture [1]. In healthy tendon, a typical hierarchical arrangement of parallel collagen fibrils and fibers forms a tendon unit, which in an unloaded state adopts a crimp-type/wavy configuration [2]. Tendon fibroblasts, i.e. tenocytes, contribute to tissue homeostasis when exposed to this highly ordered collagen extracellular matrix (ECM) by producing ECM proteins and thus collagen assembly and turnover [3], [4]. However, due to its low cellularity and hypovascularity, tendons possess a low regenerative capacity that results in poor and slow healing [5]–[7]. Injury or tissue damage often results in an increased ratio of type III collagen to type I collagen and additional deposition of glycosaminoglycans (GAG) [1]. From a structural point of view, the organization of collagen fibers change from highly anisotropic to more isotropic and at the micro-level, they become more angulated and the number of small-diameter collagen fibers is increased [8]. At the cellular level, tenocytes become more stellate-shaped [1], [8]–[10] and the expression of chondrogenic genes such as *Col2a1 and Aggrecan* increases [9], [11] while expression of tendon-related markers, i.e. *Tenomodulin (Tnmd)* [12] and *Scleraxis (Scx)* decreases [13].

Similar to *in vivo*, changes in tenocyte phenotype, shape and function are observed during *in vitro* culture of tendon-derived cells on flat tissue culture plastic, which thus lacks the tendon ECM niche. On flat tissue culture polystyrene, tenocytes quickly lose their elongated shape and decrease the expression levels of *Scx* [14]. Yao *et al*., reported that the amount of collagen type I (COL1) and decorin (DCN) protein expression levels decreased significantly from passage 0 to passage 8 during *in vitro* culture [15]. Mazzocca *et al*., showed that in addition to *Scx, Dcn* and *Col1*, gene expression levels of other tenocyte markers, i.e. Tenascin-C (*Tn-C*) and *Tnmd*, decline from passage 0 to passage 6 [16]. This phenomenon is referred to as dedifferentiation, and is also a well-known phenomenon observed in chondrocytes [17]. These results point towards the lack of a healthy tendon niche, i.e. biochemical and topographical changes, driving this loss of tenocyte phenotype, in analogy to changes in tissue organization *in vivo* that similarly affect tenogenicity.

Various strategies were used to re-differentiate the dedifferentiated tenocytes to rescue their tenocyte phenotype *in vitro*. Supplementing culture medium with various growth factors such as VEGF [18], IGF [19] and GDF5 [20] increases expression of tenocyte marker genes including *Scx* and *Tnmd*. Additionally, serum-deprivation, i.e. depletion of medium-associated growth factors rescued the lost tenocyte phenotype [21]. In addition, the small molecule tazaratone, which targets the retinoic acid receptor, preserves tenocyte phenotype via scleraxis [22]. Manipulating tenocyte shape has also been used as an approach to re-differentiate dedifferentiated tenocytes, and one such approach that pushes cells into an elongated shape is by exposing cells to a topographical cue, i.e. a substrate composed of anisotropic fibers [23]–[32]. For instance, Kishore *et al*., used aligned collagen threads to mimic the packing density, alignment and strength of native tendon, and observed an increase in gene expression of *Scx, Tnmd* and *Tn-C* compared to the randomly organized fibers [27]. Similarly, Younesi *et al*., fabricated 3D-bio textiles from collagen and observed that compared to random fibers, expression of *Col1a1* and *Tnmd* increased more than 6- and 11-fold, respectively [33]. Tu *et al*., promoted tenogenic differentiation *in vitro* and tendon regeneration *in vivo* by aligned electrospun fibers derived from tendon ECM [29]. An increase in the expression of tenogenic marker genes, including *Scx, Dcn* and Biglycan (*Bgn*), was observed both *in vivo* and *in vitro* [29]. We previously reported that tendon-derived cells populating a reconstituted collagen tissue, ranging from uniaxially constrained with spindle-shaped cells to biaxially constrained with stellate-shaped cells, display the strongest functional remodeling capacity and tenocyte phenotype when adopting a spindle-shaped morphology [34]. In another approach, we took the cell shape concept one step further by using a micro-topographical screening platform with a large amount of unique micro-topographies for identifying feature-characteristics associated with Scleraxis expression, and revealed a strong correlation of specific cell shapes, i.e. cell and nuclear area with Scleraxis levels [14]. These studies underline the impact of stimulating the tenocyte phenotype by manipulating the cell shape.

Based on the above, we hypothesized that the topographical cues present in the tenocyte niche contribute to phenotypic maintenance of tenocytes. We tested this by exploiting an imprinting method to replicate the native tendon topography to polystyrene. Results of this study showed the importance of the tendon topography, pushing cells in a spindle-shaped morphology, in the maintenance of the tenocyte phenotype *in vitro*, which can shed a light on further understanding of tendinopathy.

## Results

### Tendon-derived cells rapidly change morphology and phenotype on a flat substrate

Flat polystyrene surfaces were used to assess the influence of passage number on tenocyte shape and tenocyte phenotype. Therefore, passage one (P1) rat tenocytes were cultured and subsequently passaged four times (Fig. 1). We first investigated the changes in cell and nuclear shape parameters including area, compactness, solidity and aspect ratio and changes in gene expression levels of tendon-related genes. Aspect ratio is calculated by the division of the length of the major axis of by the minor axis; so, the higher the value, the more elongated the cells. Compactness is correlated with elongation; high compactness values indicate more elongated cells. Solidity is related with cell branching and the closer its value to 1, the more solid the cells. Results showed a slight increase in cell area, i.e. cell size, with passage number (Fig. 1B). Cell aspect ratio and compactness significantly decreased after P1, yet solidity remained the same in all groups (Fig. 1B). The increase in the nucleus area was statistically significant between P1 and P3, P4 but remained at similar values at P2 and P5 (Fig. 1B). Nucleus aspect ratio significantly reduced after P2, yet compactness and solidity remained unchanged between passages (Fig. 1B, supplementary file 1 containing raw data). Overall, our data indicates that with passaging of tenocytes to P4 on flat tissue culture plastic, they undergo shape-related changes; cells and their nucleus steadily increase in size and transform from the original spindle-shaped/elongated morphology to a stellate-shaped/round morphology.

**Figure 1.**
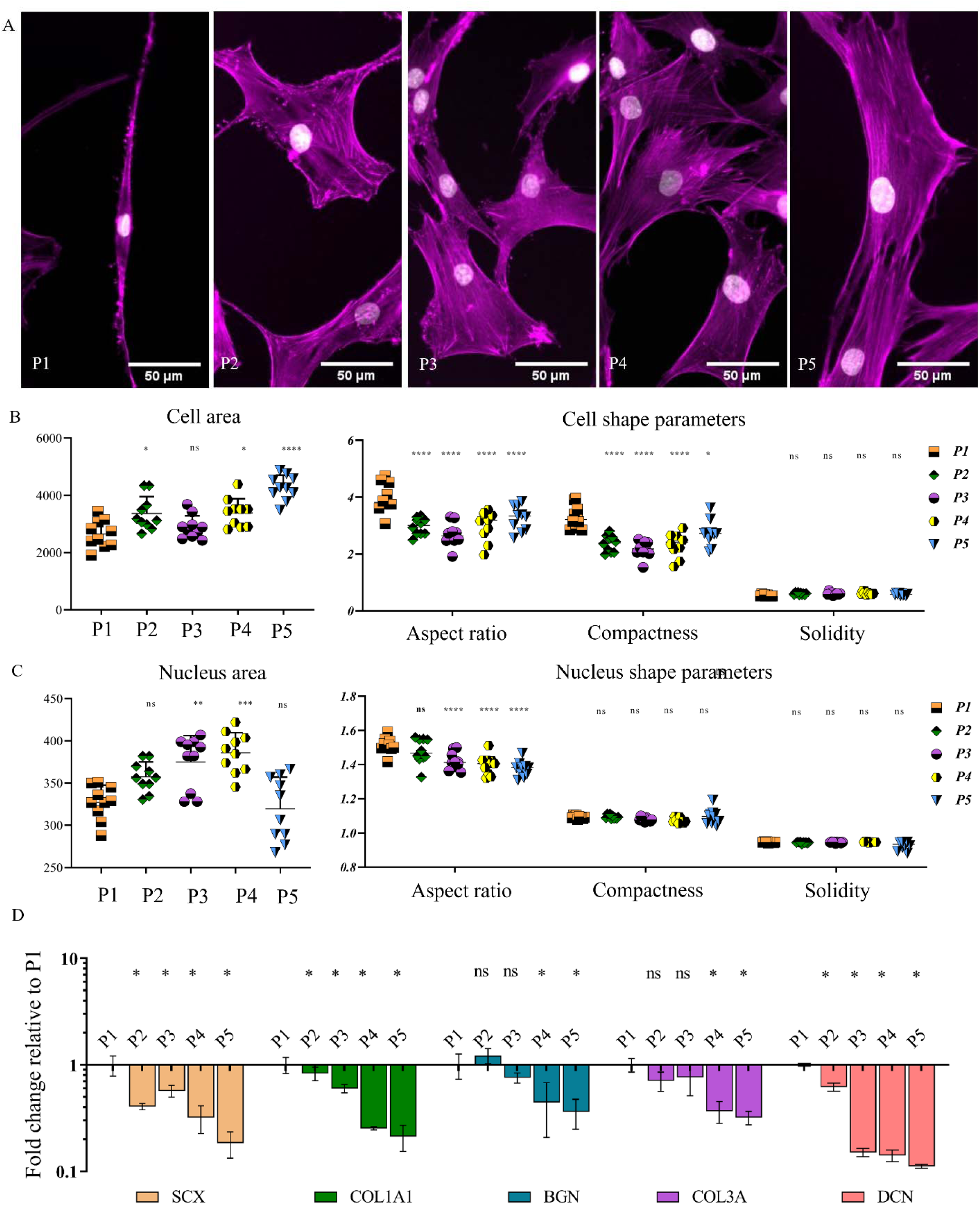
Tendon-derived cells rapidly lose their morphology and phenotype on a flat substrate. (A) Representative images of rat tenocytes stained with Phalloidin-568 to visualize F-actin filaments (Magenta-artificial color) and counterstained with DAPI to visualize the nucleus (Cyan-artificial color) cultured on flat surfaces from passage 1 (P1) until passage 5 (P5). (B) Changes in cell area, aspect ratio, compactness and solidity from P1 to P5 (*P < 0.05, **P < 0.01,***P < 0.005, ****P < 0.0001). (C) Changes in nucleus area, aspect ratio and solidity from P1 to P5 (**P < 0.01,***P < 0.005, ****P < 0.0001). (D) Expression of tenogenic marker genes Scleraxis (*Scx*), Collagen type 1 (*Col1a1*), Collagen type three (*Col3*), Decorin (*Dcn*) and Biglycan (*Bgn)* from P1 towards P5 all decreased with passage number. A log_10_ transformation of expression fold changes is used to visualize the differences in gene expression. Scale bars represent 100 μm. (Error bars represent 95% confidence intervals, *P < 0.05,**P < 0.01). For all experiments, n = 3.

To further evaluate the loss in tenocyte phenotype, gene expression levels of *Scx, Col1a1, Col3, Dcn* and *Bgn* were measured (Fig. 1D)). *Scx*, which is a tenogenic transcription factor [35], showed a significant decrease after passage one. A similar reduction was observed for the expression of *Col1a1*, which is the most abundant protein in the tendon extracellular matrix (ECM), and *Col3* [36]. Furthermore, we measured the expression of genes coding for non-collagenous matrix proteins *Dcn* and *Bgn* that bind to collagen, which displayed a decreasing trend from P1 to P5 [37]. With these results, we confirmed that tenocytes dedifferentiate by rapidly losing their morphological characteristics and expression of tenocyte marker genes on a flat surface during *in vitro* sub-culture.

### The native tendon imprint as cellular niche on polystyrene

After establishing that with passaging, tenocytes lose their phenotype on flat polystyrene, the effect of the tendon surface topography, as an isolated effect, was assessed. To this end, we used the soft embossing method to imprint a native tendon topography onto polystyrene (Fig. 2). The imprinting protocol involved polymerization of PDMS using a 10:1 ratio of monomers and curing agent on a native tendon (Fig.2A, 3), and embossing the PMDS negative imprint on polystyrene (Fig.2A,6). The imprinted material was imaged using scanning electron microscopy (SEM) (Fig.2B and supplementary video 1), atomic force microscopy (Fig.2C) and profilometry (Fig.2D and supplementary video 2). SEM and profilometer images reveal that native tendon can indeed be imprinted onto polystyrene, and the polystyrene imprint possesses a tendon-like topography. AFM analysis shows that surface topographies on tendon imprints can reach a depth of 5μm.

**Figure 2.**
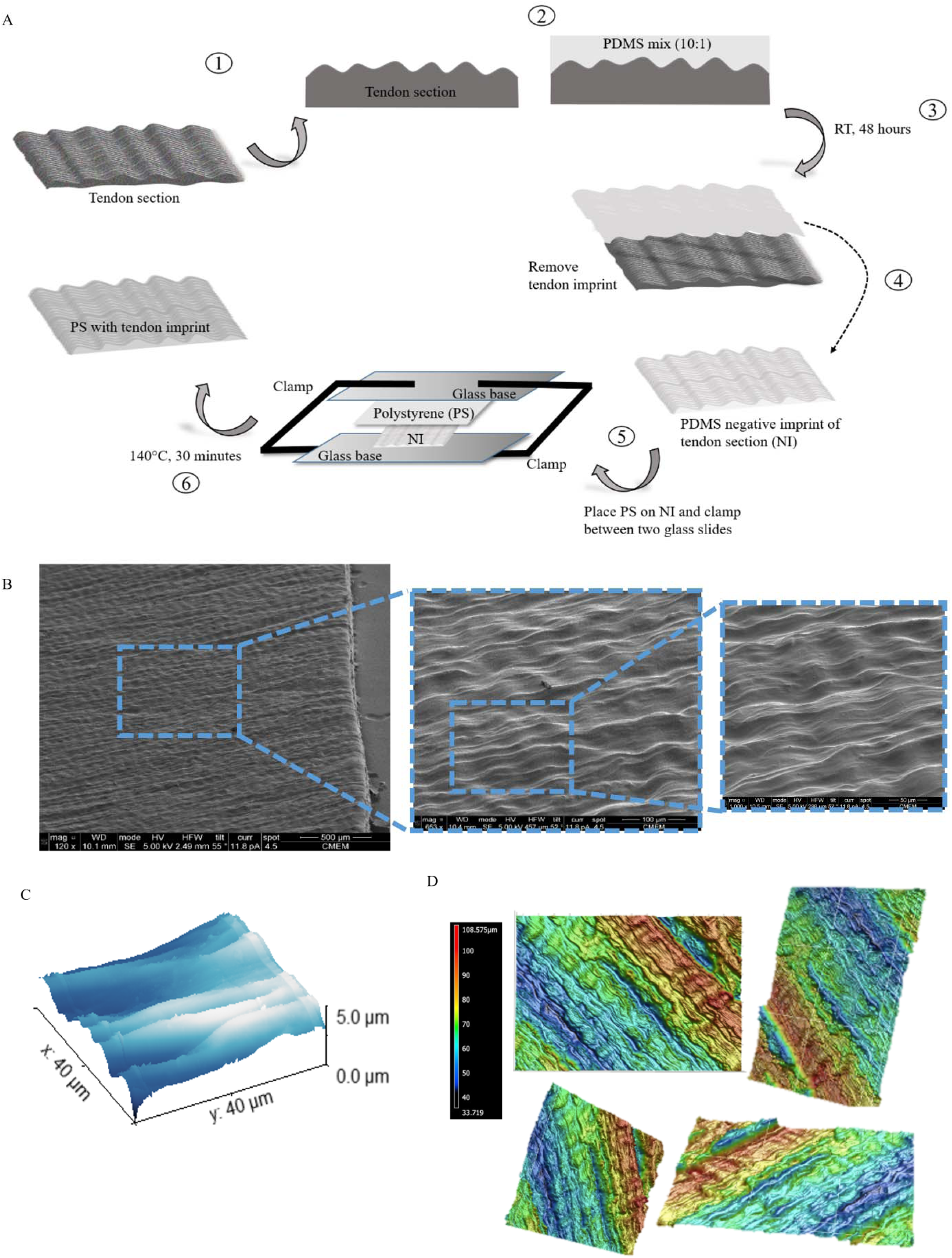
The native tendon imprinted as a cellular niche on polystyrene. (A) Procedure to develop a native tendon imprint, using an embossing technique of a native tendon tissue into polystyrene. (B) SEM image of tendon imprint with lower (left) and higher (right) magnification. The red dotted box represents a randomly selected region on the imprint. (C) Representative AFM image of tendon imprint showing that the imprint can reach a depth of 5μm height. (D) Representative image of profilometer images of tendon imprint, depicted from different angles.

### Late passage tenocytes adapt early passage tenocyte morphology on tendon imprints

Next, the morphological response of tenocytes to the tendon imprint was assessed (Fig.3). Flat control surfaces were prepared by imprinting a flat PDMS mold to the polystyrene sheets. Rat tenocytes (P4) were seeded on flat and imprint surfaces and stained with phalloidin to visualize F-actin (grey) and DAPI to visualize nuclei (blue). On the flat surface, (Fig.3A&B, top panel) a spread morphology was observed, contrary to tendon imprints, where cells on each replicate displayed an elongated shape (Fig.3A&B, bottom panel).

**Figure 3.**
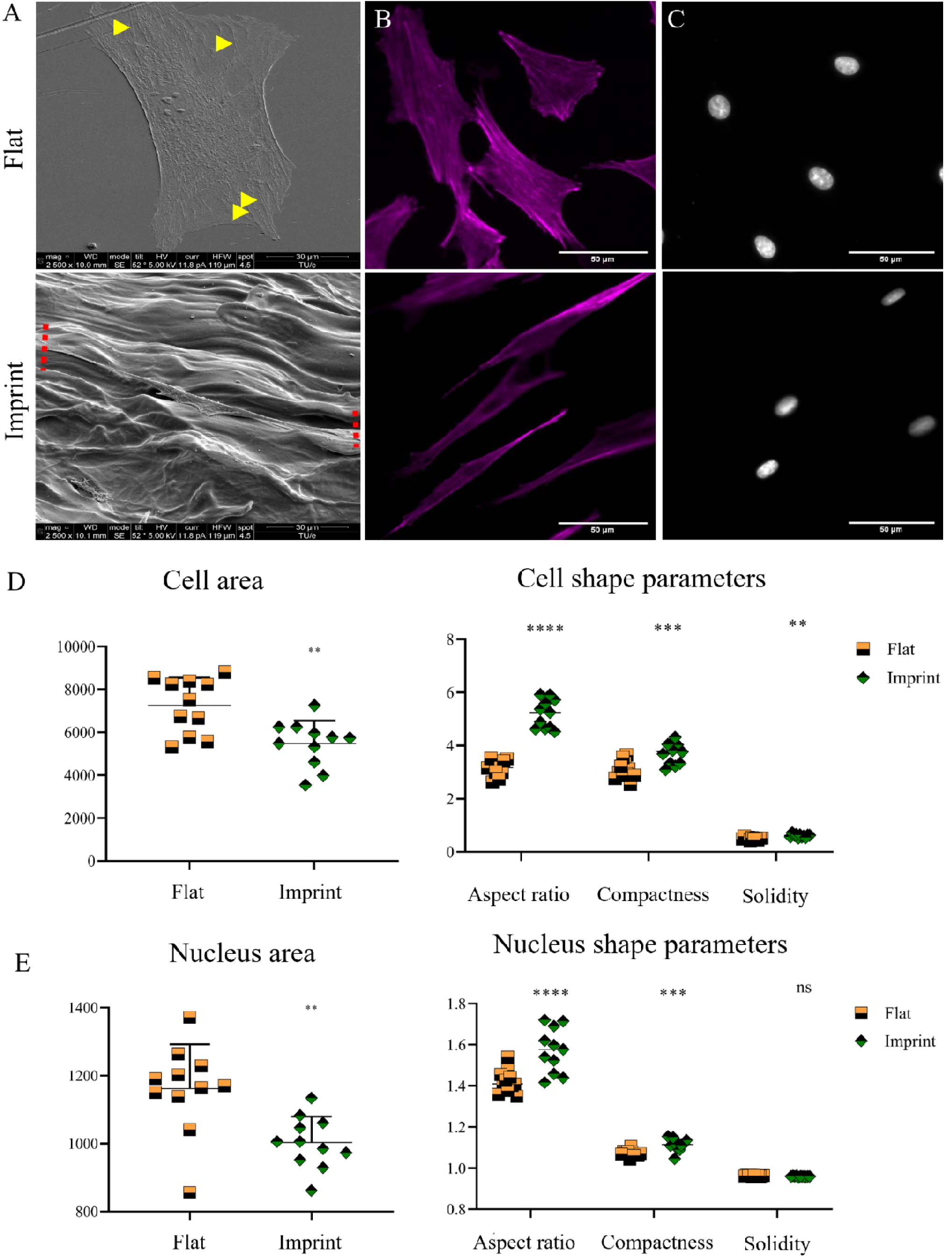
Tendon imprint topography manipulates tenocytes to their natural morphology. Tendon imprint topography pushes tenocytes to their natural morphology (A) Representative images of tenocytes cultured on the flat surface (left) and four replicates of the tendon of tendon imprints (right). Top images belong to SEM imaging, bottom images belong to fluorescent staining (Phalloidin in Gray, artificial color. DAPI in blue). Scale bars represent 100 μm. (B) SEM images of tenocytes cultured on a flat surface (top) and tendon imprint (bottom). Yellow arrows represent stress fibers on the tenocytes on the flat surface. Red dotted lines represent the major axis boundaries of tenocytes. Scale bars represent 30 μm. (C) Representative images of cell shape (magenta, artificial color) of rat tenocytes cultured on flat (top) and tendon imprint (bottom). Scale bars represent 10 μm. (D) Representative images of nucleus shape (grey, artificial color) of rat tenocytes cultured on flat (top) and tendon imprint (bottom). Scale bars represent 10 μm. (E) Cell shape parameters are significantly different between both flat and imprint topographies, except for nuclear solidity (* p<0.05). For all experiments, n = 3.

We further investigated the effect of tendon topography on cell and nuclear shape parameters including aspect ratio, area, solidity and compactness (Fig.3D&E). We first visualized the difference in cell shape. Rat tenocytes were exposed to either a flat surface or tendon imprint topography and subsequently SEM imaging was performed (Fig.3A). On the flat surface, we observed that tenocytes are more spread and stress fibers were observed (yellow arrows) (Fig.3A-top panel). On tendon imprints, rat tenocytes were highly elongated (red arrow) (Fig.3A-bottom panel). We observed similar results when we stained tenocytes with phalloidin to visualize the actin cytoskeleton (Fig.3B) and DAPI to visualize nuclei (Fig.3C). Cell and nuclear area were significantly smaller on tendon imprints compared to flat (Fig.3E&F). Additionally, cellular and nuclear aspect ratio and compactness were significantly higher on tendon imprints, compared to flat (Fig.3E). Cell solidity on imprints (0.61 ± 0.05) was significantly higher and closer to 1, compared to the flat surface (0.51 ± 0.04); indicating that cell branching was less pronounced on tendon imprint. Nuclear solidity was not significantly different for both topographies. Therefore, results indicate that tendon imprints induce an elongated shape and reduces cytoskeletal branching, resembling the early passage tenocytes on flat polystyrene.

### Tendon imprint topography decreases tenocyte proliferation

Surface topography has been identified as a regulator of cell proliferation [38] [39]. Therefore, the effect of tendon imprint topography on tenocyte proliferation compared to the flat surface was assessed. Rat tenocytes were thus cultured on a flat surface and tendon imprint, and after 24 hours the proportion of cells in the S-phase of the cell cycle (pink-purple) to all cells (blue) was measured by EdU labelling (Fig.4A&B). On tendon imprints, 37% of the tenocytes were EdU positive whereas on a flat surface, 63% of the cells were EdU positive. This indicates that tendon topography relates to a less proliferative tenocyte state.

### Expression of SCX correlates with cellular and nuclear features

Tendon imprints significantly affect tenocyte cell and nucleus shape. Next, the influence of tendon imprint topography on tenocyte marker gene expression was assessed (Fig.5). Therefore, rat tenocytes at P4 were cultured on tendon imprints and flat surfaces for 24 hours and the amount of SCX was measured by immunocytochemistry and subsequent quantification (Fig.5A). SCX levels were significantly higher expressed on tendon imprint after 24 hours (Fig.5B), which was subsequently correlated with cell shape, using a Spearmen correlation calculation. A positive and significant correlation was detected of SCX level with cell compactness (r_s_= 0.71), nucleus compactness (r_s_= 0.71), cell eccentricity (r_s_= 0.82) and nucleus eccentricity (r_s_= 0.78) (Fig. 5C&D, supplementary file 2 containing raw data). In addition, a negative and significant correlation was detected of SCX level with cell area (r_s_= −0.58) and nucleus area (r_s_= −0.70) (Fig. 5C&D). SCX level thus correlates with elongated cell and nucleus shape, driven by tendon surface topography.

**Figure 4.**
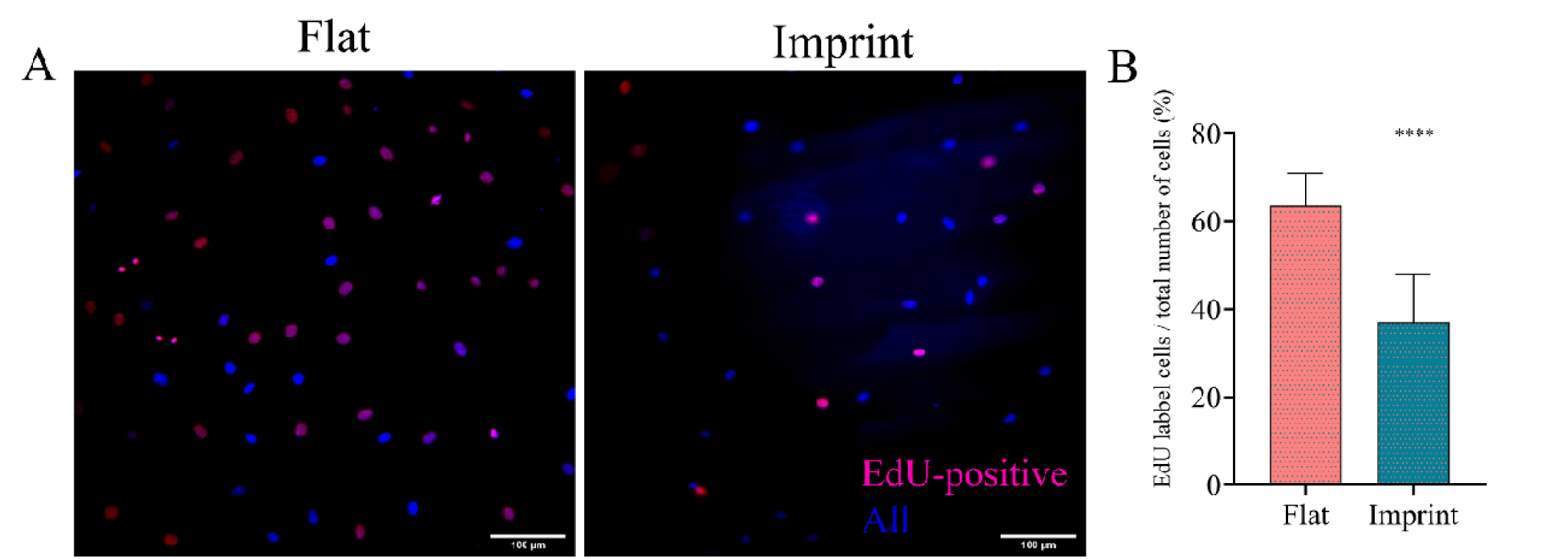
Tendon imprint topography lower cell proliferation capacity. (A) Representative images of EdU staining in which proliferating cells are labelled in pink-purple color and Hoechst is used as a counterstain (in blue color). (B) Quantification of EdU positive tenocytes on the flat surface and tendon imprints. On the flat surface, 63% of the cells was EdU positive, whereas on tendon imprints it was only 37%. Scale bars represent 100 μm. (Error bars represent 95% confidence intervals, * p<0.05). For all experiments, n = 3.

**Figure 5.**
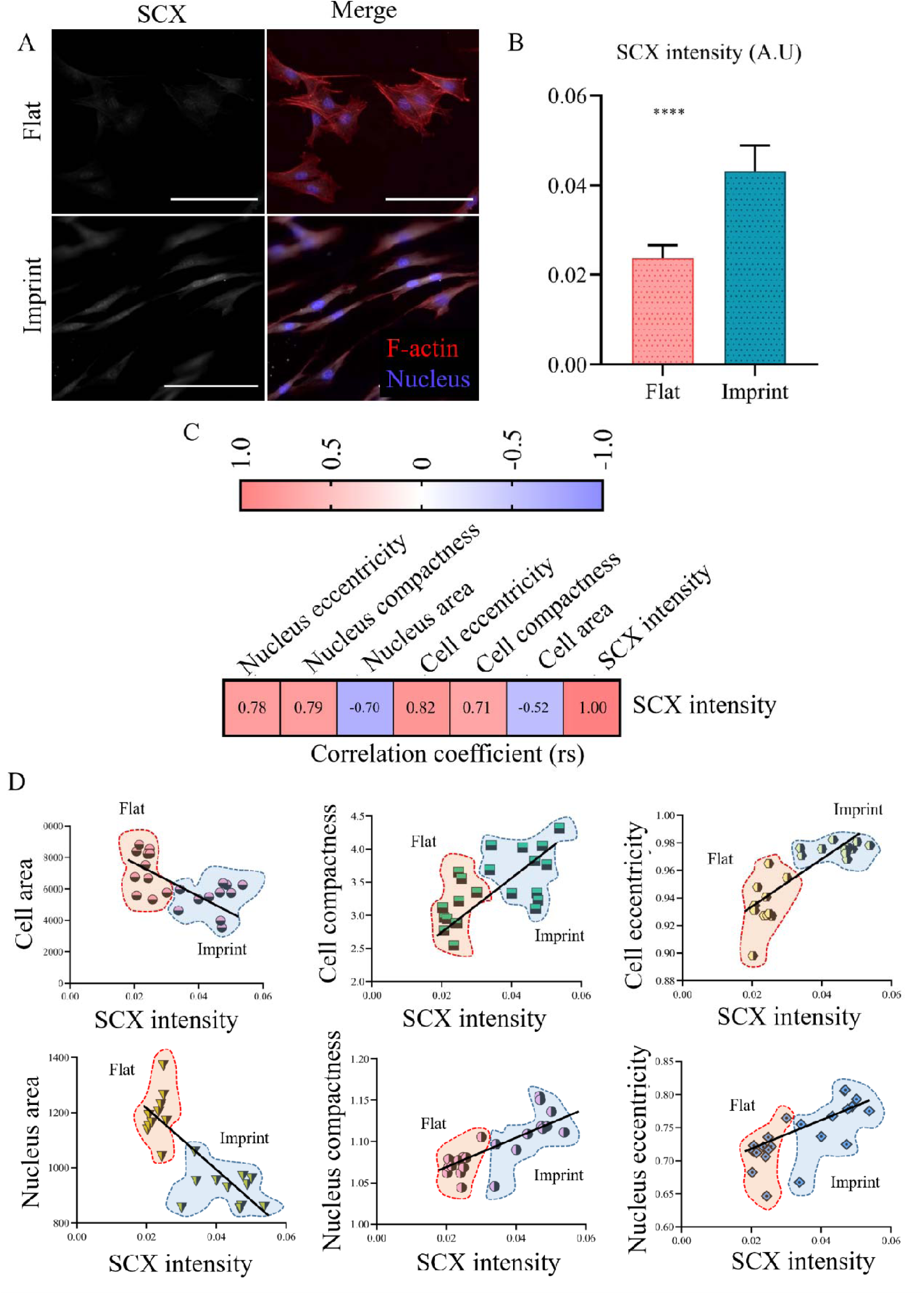
Expression of SCX correlates with cellular and nuclear features. Tenocyte phenotype is maintained by tendon imprints (A) Representative images of rat tenocytes cultured for 24 hours on a flat surface and a tendon imprint and stained for SCX. (B) Quantification of SCX level in tenocytes on flat surface and tendon imprint. (C) Levels of SCX correlates with cellular and nuclear features Spearmen correlation values (r_s_) between SCX levels and eccentricity, compactness and area of cell and nucleus. (D) X-Y graphs of cell and nuclear area, compactness and eccentricity correlated with SCX level. Each data point represents the median value of individual intensities of all cells measured in a single image. Area indicated in red and blue belong to cells on flat and imprint surfaces, respectively. For visual purposes, a trend line was added derived from a simple linear regression calculation.

### Tenogenic gene expression is maintained on tendon imprints

After showing that native tendon topography can increase SCX level on the short term, we investigated whether tenocyte phenotype can be maintained for 7 days of culture. (Fig. 6). Therefore, P4 tenocytes were cultured on flat surfaces and imprints for 7 days and gene expression levels of *Scx, Col1a1, Sox9, Dcn*, and *Runx2* were measured on day 1 (D1), day 3 (D3) and day 7 (D7). On the flat surface, expression of tendon-related genes *Scx, Dcn* and *Col1a1* decreased over 7 days of culturing, indicating ongoing dedifferentiation (Fig. 6A-C). Expression of *Sox9*, a chondrogenic transcription factor that is also expressed at the tendon-bone junction, also decreases on the flat surface (Fig. 6E). Lastly, *Runx2* gene expression, an osteogenic-associated transcription factor, significantly increased on the flat surface. Contrary, on tendon imprints, gene expression levels of *Scx, Dcn, Col1a1* and *Sox9* increased at D1 and subsequently remained stable over the 7 days, indicating that tendon imprints can both induce a slight increase in tenogenic gene expression and prevent the dedifferentiation that occurs on the flat surface (Fig. 6A-C, E). Lastly, gene expression of *Runx2* did not increase on the tendon imprints.

**Figure 6.**
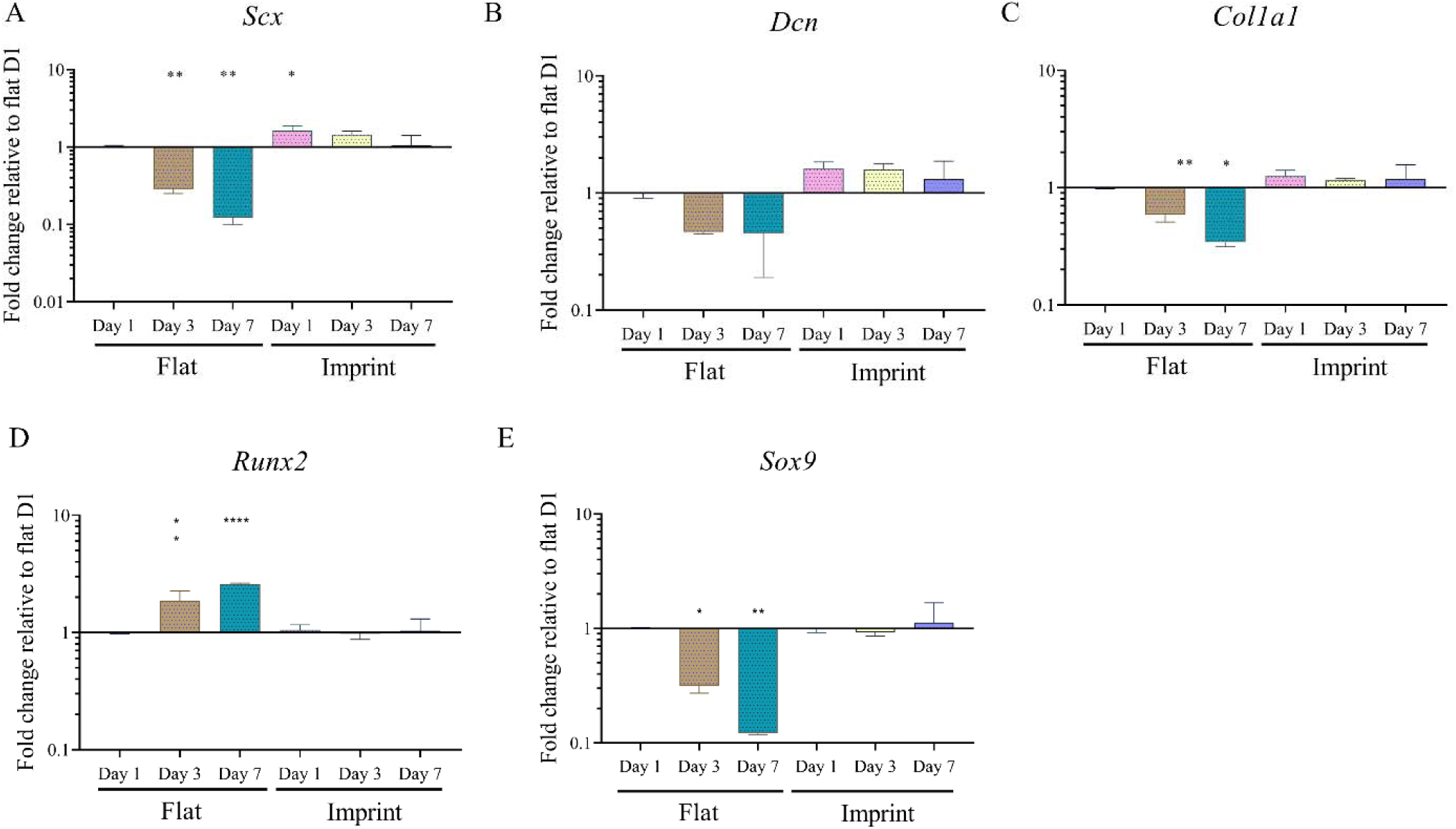
Tenocyte phenotype is maintained on tendon imprint topography. Gene expression levels of (A) *Scx*, (B) *Dcn*, (C) *Col1a1*, (D) *Runx2* and (E) *Sox9* of tenocytes cultured on flat surfaces and tendon imprints over 7 days of culture. On a flat surface, expression of *Scx, Dcn, Col1a1, Sox9* decrease whereas *Runx2* increases. Contrary, tendon imprints preserved tenocyte phenotype. A log_10_ transformation of the expression fold changes is used to visualize the differences in gene expression. (Error bars represent 95% confidence intervals, *P < 0.05,**P < 0.01). For all experiments, n = 3.

## Discussion

Surface topography of various organs and tissues in nature give a series of evolutionary advantages and provide specialized functions. They provide cells with signals to differentiate, migrate, proliferate or maintain their phenotype. The surface topography of tendon tissue is also considered to provide such strong cues. However, whether the tendon topography alone, devoid of other cues (e.g. ligand type and density, changing mechanical properties, etc), provides a strong cellular cue remained to be explored. Therefore, the soft embossing method was exploited to imprint native tendon on a polystyrene surface to study the effect of tendon surface topography on cell shape, proliferation and phenotype, when compared to flat polystyrene surfaces. Results of this study highlight that: 1) native tendon topography can be imprinted onto a polystyrene surface to conduct tendon-related studies *in vitro*, 2) flat polystyrene surfaces drastically alter tenogenic cell shape and phenotype, 3) cellular proliferation is decreased on imprint topographies, compared to flat surfaces, 4) tenocyte phenotype is maintained on tendon imprint topographies and 5) remarkably, Scleraxis expression, a key tendon marker, beautifully correlates with specific cell and nuclear shape parameters. Concordantly, results of this study imply that imprinting the natural tendon surface on materials, such as polystyrene, provides a tool to control tenocyte cell shape and maintenance of the tenocyte phenotype, as summarized in Fig. 7.

**Figure 7.**
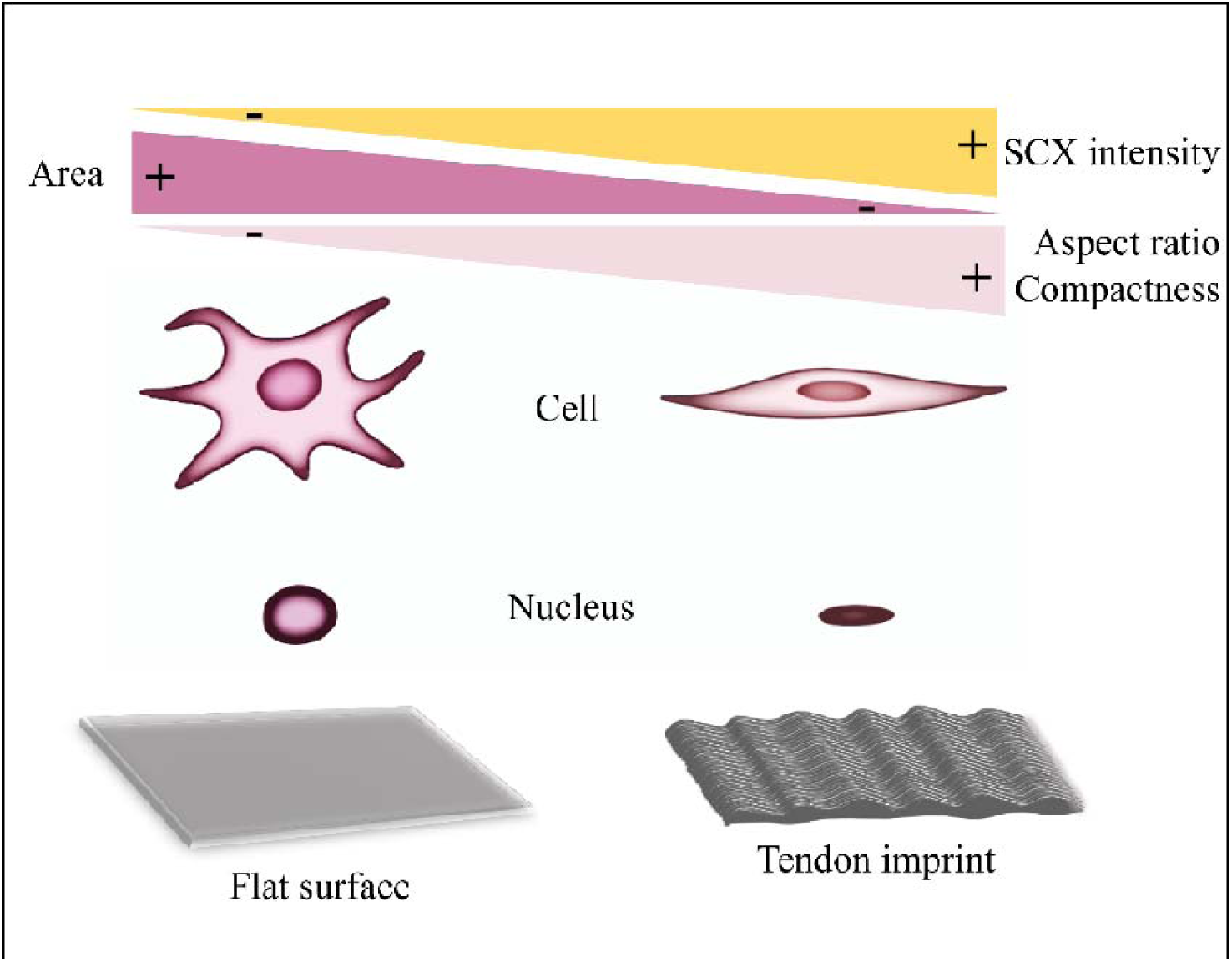
Summary of the results. Tenocytes on imprint surfaces display a lower cell area and higher aspect ratio, compactness and solidity, which results in increased SCX intensity.

Drastic changes in cell shape of freshly isolated or early passage cells upon culturing on tissue culture plastic were shown for a wide variety of cell types, including tenocytes [15], [14], [40]– [45]. Vermeulen *et al.* and Yao *et al.* showed that the elongated shape of tenocytes at P0 transforms to stellate-shaped at P1 [15], [14], which is in agreement with the changes observed in cell and nucleus shape of the current study. Similarly, our observation on the decrease in tenogenic marker genes corroborate with previously published reports [14], [15], [16], indicative for a dedifferentiation process. Due to the rather limited availability of the early passage cells (P0 or P1), dedifferentiation has become an obstacle for *in vitro* research and associated interpretation of acquired data of many cell types. This gives rise to the interesting concept of re-differentiation in order to promote/restore the native phenotype.

The interplay between cell shape and cell fate (e.g. differentiation, proliferation, apoptosis etc.) is an established concept [55] [17]. Following that concept, because healthy tenocytes *in vivo* adopt an elongated morphology, i.e. the nucleus and cell body, which are aligned between collagen fibers, re-differentiation research has focused on pushing cells towards similar morphologies. Among these approaches, cells have been exposed to anisotropic fibers [24], [27], [31], [32], [46]–[52] or nano/micro-grooves [14], [26], [38], [53]. For instance, Zhu *et al.*, reported that dedifferentiated tenocytes become elongated upon culturing on microgrooves, which partially reversed the dedifferentiation process [26]. A similar study by Schoenenberger *et al.*, showed that tendon fibroblasts displayed a higher expression of the tendon-related marker *Mohawk* (MKX), COL, BGN and DCN on aligned PCL fibers, compared to random fibers [54]. Results of the current study supports the concept that enforcing a spindle-shaped tenocyte morphology is beneficial for maintaining/restoring their native phenotype. Specifically, our results showed that elongated cell shape, i.e. high cell aspect ratio, eccentricity and compactness, is correlated with SCX levels. However, a degree of heterogeneity in cell phenotype, i.e. shape parameters and SCX levels, was observed especially for tendon imprints. Explanations for this may relate to the imprinting technique that potentially disturbs native architecture locally, or the fact that cells on imprints are exposed to a 2D surface with more freedom to adopt a preferential shape, rather than the strongly confined 3D in vivo environment. Despite that, the expression of tenocyte marker genes *Scx, Dcn* and *Col1a1* was still shown to be stabilized on tendon imprints, contrary to a decay observed on flat surfaces.

The observation in the current study that osteogenesis (*Runx2*) increased on flat surfaces, compared to imprints, was also observed in another study, using osteocalcin and alkaline phosphatase as osteogenic marker genes [31]. Possibly, this relates to the increased proliferation rate on flat surfaces resulting in cell confluency, a phenomenon that was previously linked to increased *Runx2* gene expression [14], [24], [56]. Although tenocytes are the most abundant cell type in tendon tissue, depending on the location of the cells within the tendon, a *Sox9* positive subpopulation of tenocytes exists that is located at the junction between bone and tendon/ligament [57], [58]. The expression of *Sox9* was previously shown to remain stable when tenocytes were cultured, either cyclically stretched or left unstretched, on microgrooved substrates, which induces an elongated morphology [59]. Therefore, the naturally present *Sox9* gene expression level in elongated tenocytes, which is stabilized on imprint surfaces, is lost on flat surfaces since the differentiation process is initiated. Suggestively, different topographical cues present *in vivo* in the tendon towards the enthesis, contribute to differences in phenotype of locally present cells.

Healthy tendon comprises of relatively quiescent tenocytes, whereas proliferation increases during the remodeling phase of tendon during healing, during which the strong tissue anisotropy is absent [60]. In analogy, tendon imprints resulted in suppression of proliferation, when compared to flat surfaces. Similarly, proliferation and the differentiation capacity towards the tenogenic lineage of human mesenchymal stem cells is also lower on aligned collagen fibers or anisotropic topographies, compared to randomly oriented substrates [61][38]. One explanation for different proliferation rates on different topographies is a difference in cell shape and alignment of stress fibers, as previously shown [62]. In elongated smooth muscle cells (SMCs), DNA synthesis was decreased, indicating that the directed cell spreading can be positively linked with DNA synthesis and thus cell proliferation [63]. A possible underlying mechanism for this could relate to *NOR-1*, a gene involved in SMCs proliferation. *NOR-1* expression is among others modulated by protein kinase C (PKC). In elongated cells, the activity of protein kinase C was shown to be decreased, which was associated with a decreased *NOR-1* expression and thus ultimately decreased cell proliferation [62]. Considering the similarities in shape influences between SMCs and tenocytes, this could be a possible explanation. Another, or additional, explanation is that changes in cell proliferation are linked to surface-topography-induced mechanotransduction. Surface topography was shown to alter integrin activation and focal adhesion dynamics, which affects cell proliferation [64], [65]. Flat surfaces results in specific cell spreading and integrin binding, resulting in focal adhesion kinase phosphorylation and activation, which in turn activates the ERK pathway, cyclin D1, and finally the G_1_-S cell cycle transition [66]. However, activation of integrins α5 and α6 have been shown to create the opposite effect [67]. This indicates that the exact mechanism by which tenocyte proliferation is coupled to cell shape, surface topography, mechanotransduction pathways and specific integrin recruitment on tendon imprints should further be explored.

The use of tendon imprints in research comes has several advantages. Firstly, the production of tendon imprints requires only PDMS, polystyrene and tools to perform soft embossing; therefore, upscaling the production can be cheap and fast. Secondly, they bear the potential to be used as a model system to study the molecular mechanism behind the dedifferentiation and redifferentiation process, including the role of surface topographies on the activation of mechanotransduction pathways. Thirdly, the imprinting technology is facile enough that it can be applied to any polymer that does not degrade above 140°C. Therefore, in addition to the influence of healthy tendon surface topography, the effect of other polymers on tenocyte phenotype can be investigated. Finally, the technique also allows imprinting the pathogenic or damaged tendon topography on polystyrene; therefore, the role of altered surface topography on tenocyte phenotype can be further explored *in vitro*.

## Conclusion

This study shows that tenocyte cell shape and expression of tenogenic marker genes drastically change during *in vitro* subculture, which leads to decreased tenogenic characteristics. Furthermore, we demonstrated that tendon imprints, which carry the topological cues from the native tendon, led to an elongated cell shape and resulted maintenance in the expression of tenogenic-associated markers genes, which is positively correlated with elongated cell and nuclear shape parameters, i.e. aspect ratio, compactness and eccentricity. Overall, results of this study support the concept of cell shape-to-phenotype in tenocytes and stresses the role of surface topography on tenocyte phenotype.

## Data availability statement

All data is available upon request.

## Acknowledgements

This research has received funding from the European Union’s Horizon 2020 research and innovation programme under the Marie Sklodowska-Curie grant agreement No 676338.

## Disclosure statement

Authors do not have any financial interests to declare.

## Appendix A. Supplementary data

### Materials and Method

#### Tendon tissue imprinting

Fresh porcine Achilles tendons were obtained from crossbreeds of Great Yorkshire and Dutch land pigs aged between 6-8 months old and between 85-95 kg of weight, supplied by a local slaughterhouse (Compaxo Meat B.V, the Netherlands). Muscle, fat, bone-like tissues, synovial sheath, and paratenon were aseptically dissected and the remaining tendon was cut into 1cm^3^ blocks and stored at −80 °C. Frozen tendons were embedded in a mixture of polyvinyl alcohol and polyethene glycol (OCT, Sakura) and fixed to the cutting base plate of a cryotome (Leica CM1950) after which longitudinal sections were cut with a thickness of 300 μm. Sections were washed with PBS and stored at −80 °C until use.

Tendon sections were thawed at room temperature for 30 minutes and placed in a 6-Well plate. Polydimethylsiloxane (PDMS, Dow Corning Sylgard 184, 4019862) was mixed with curing agent at a ratio of 10:1 (w/w) and mixed vigorously. In order to remove the air bubbles, the mixture was centrifuged at 3000 G for 10 minutes and poured on tendon sections. They were allowed to cure for 48 hours at room temperature on a stable flat surface. Then, the tendon section was peeled off from the PDMS resulting in a negative imprint of a tendon on PDMS. In between two glass slides, a negative imprint of PDMS was placed onto a polystyrene film (PS, GoodFellow) and pressed together with clamps and incubated at 140°C for 30 minutes. Flat surfaces were prepared by performing the same embossing method but using a flat PDMD mold. In order to allow cell attachment, polystyrene surfaces were oxygen plasma treated for 45 s at 75 mTor, 50 sccm O2, and 50□W.

#### Isolation of rat tenocytes

Rat tenocytes were isolated from the Achilles tendon of 23 weeks old Cyp1a2ren strain rats, after euthanization due their surplus status from the breeding program. Briefly, tendons were cut into small pieces and digested in a buffer containing 3mg/ml collagenase type II (Worthington Biochemical), 4mg/ml dispase II (Sigma-Aldrich) and 100□U/ml Penicillin/Streptomycin (Thermo Fisher Scientific) for 4 hours at 37°C in a humidified tissue culture chamber with 5% CO2. Then, the suspension was passed through a 70 mm cell strainer (Life sciences) to obtain a cell-only suspension. The cell suspension was centrifuged at 300 G for 5 minutes and re-suspended in Dulbecco’s modified Eagle’s medium (DMEM, Sigma-Aldrich) supplemented with 10% fetal bovine serum (FBS), 100□U/ml penicillin/streptomycin. Cells were cultured in T-25 flasks until 70% confluency.

#### Sterilization of tendon imprints and cell culture

Sterilization of flat and tendon imprints was performed by incubating the materials in 70% ethanol for 1 hour, and remaining ethanol was air-dried under sterile conditions. Next, samples were incubated with sterile PBS for 30 minutes at 37°C and subsequently with culture medium for 30 minutes at 37°C before use.

Rat tenocytes were seeded on the surfaces at a density of 5000 cells/cm^2^ in Dulbecco’s modified Eagle’s medium (DMEM, Sigma-Aldrich) supplemented with 10% fetal bovine serum (FBS) (Sigma-Aldrich) and 100□U/ml penicillin/streptomycin (Thermo Fisher Scientific). Cells were trypsinized once they reached 70% confluency. Tenocytes at passage 4 were used, unless stated otherwise.

#### RNA isolation and quantitative PCR (RT-qPCR)

Total RNA from each sample was isolated based on the protocol described in the RNeasy Mini Kit (QIAGEN). Reverse transcription was carried out based on the protocol provided by iScript™ Select cDNA Synthesis Kit (Bio-Rad). Quantitative PCR was performed by using iQ^™^SYBR^®^ Green Supermix (Bio-Rad) by using the Bio-Rad CFX manager. Ribosomal Protein L13a (*Rpl13a)* was used as a housekeeping gene and relative expression was determined using the ΔΔCt method. Primer sequences are listed in table 1.

**Table 1.**
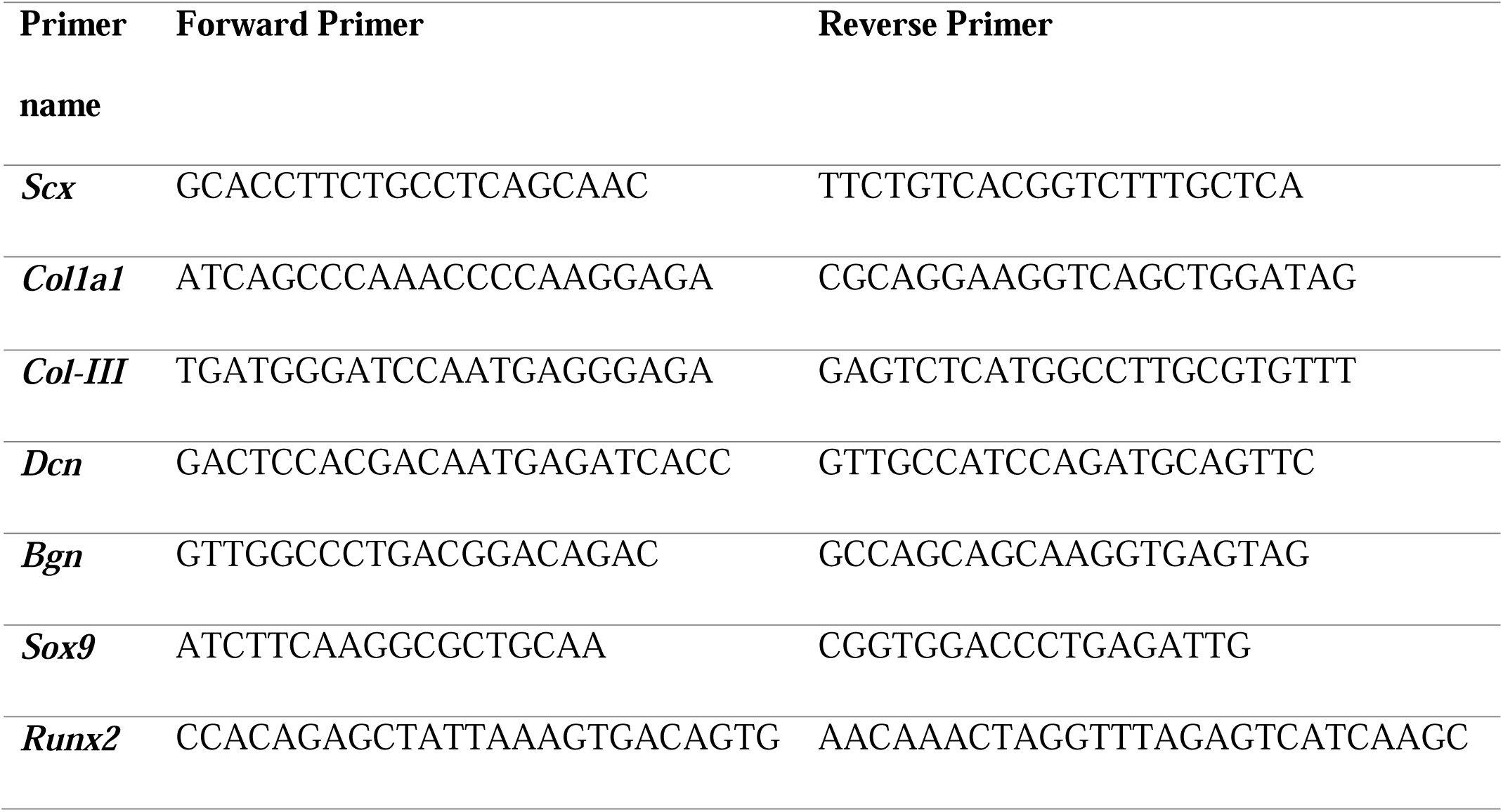
Primer sequences used in this study.

#### Immunofluorescence staining of Scleraxis expression

Cells attached to flat or tendon imprint surfaces were fixed at day 1, day 3 or day 7 after the start of culturing with 4% paraformaldehyde (PFA, ThermoFisher Scientific) at room temperature for 20 minutes and then washed with phosphate-buffered saline (PBS, Sigma-Aldrich), twice. Next, samples were permeabilized with 0.5 % (v/v) Triton X-100 in PBS for 10 minutes at room temperature. After permeabilization, cells were blocked with 1:100 horse serum in PBS for one hour at room temperature. Afterwards, samples were incubated in primary antibody for Scx (1:200; Abcam; ab58655) dissolved in 0.01% (v/v) Triton X-100 and 0.5% BSA in PBS overnight at 4°C. Next, cells were washed with 0.01% (v/v) Triton X-100 and 0.5% BSA in PBS three times and incubated with anti-rabbit secondary antibody conjugated to Alexa Fluor 647 (1:200; ThermoFisher A27040), together with Phalloidin–Tetramethylrhodamine B isothiocyanate (Phalloidin-TRITC, 1:200; ThermoFisher) in PBS with 0.01% (v/v) Triton X-100 and 0.5% BSA in PBS for 1□hour. Nuclei were stained with 4′,6-diamidino-2-phenylindole (DAPI, 1:500; Sigma-Aldrich) for 1 hour after washing. Finally, samples were mounted on glass cover slides with mounting medium (Dako, Agilent). Imaging was performed by using Leica DMi8 with a TIRF Multi Color microscope (Leica Microsystems CMS) with lasers at excitation wavelengths of 532 nm and 647 nm, for phalloidin and scleraxis respectively.

#### EdU labelling

In order to identify the proliferating cells, Click-iT™ EdU Cell Proliferation Kit (Invitrogen, C10340) for Imaging (Thermo Fisher) was used based on the manufacturer’s instructions. Briefly, tenocytes were serum-starved for 24 hours prior to EdU labelling in order to set the biological clock of the cells equally. Samples were fixed with 4% paraformaldehyde (PFA, ThermoFisher Scientific) at room temperature for 20 minutes and permeabilized with with 0.5 % (v/v) Triton X-100 in PBS for 20 minutes after 24 hours of incubation in 10□μM EdU solution. Afterwards, cells were treated with EdU reaction cocktail for 30 minutes in the dark and incubated in Hoechst for another 30 minutes. Images were taken with a Leica DMi8 with TIRF Multi Color microscope (Leica Microsystems CMS) at 20x magnification. The reported number of proliferating cells was reported as the number of EdU labelled cells/total number of cells.

#### Atomic force microscopy (AFM) and Profilometer

Tendon imprints were imaged for surface architecture by using a tapping mode atomic force microscopy (AFM; XE-100, Park Systems) by using non-contact cantilevers (PPP-NCHR, Park Systems). Data were recorded with XEP software and GWYDDION software was used to image the data. A Keyence VK-H1XM-131 at 20x magnification was used for profilometer images.

#### Scanning electron microscopy (SEM)

Samples were fixed with 2.5 % glutaraldehyde (Fisher Scientific) at room temperature for one hour. Then, they were washed with distilled water 3 times for 10 minutes, dehydrated in 25%, 50%, 75%, 90%, and 100% ethanol for 15 minutes each, and incubated in 100% ethanol for an additional 15 minutes. Next, samples were dried in Hexamethyldisilazane (HMDS) (Sigma-Aldrich) for 1 hour. Prior to imaging, samples were coated with 5 nm gold-palladium and imaged using a scanning electron microscope (SEM) (FEI Quanta 3D FEG Dual Beam).

#### Image analysis

An image analysis pipeline was created in CellProfiler version 3.19. To calculate expression levels of SCX and other shape parameters, 3-4 random images from three biological replicates was selected. Median values of all cells were per image calculated by CellProfiler and used to calculate theSpearmen correlation between median SCX intensity and cell shape parameters.

#### Statistical analysis

All statistical analyses were performed by using GraphPad Prism version 8.0 (GraphPad Software, Inc., San Diego, CA). Student’s t-test was performed to calculate statistical difference between cell and nuclear areas. One-way analysis of variance (ANOVA) were carried out to calculate the statistical difference in RT-qPCR experiments and calculate cell/nuclear shape parameters between different passages and imprint vs flat surface. Spearman correlation coefficient was calculated by GraphPad correlation calculation option. For all statistical analysis, significance set at p□<□0.05 to determine the significance between means. All quantitative data represented in this study are based on triplicated experiments.

## Notes

### Competing Interest Statement

The authors have declared no competing interest.

## References

[1] N. L. Millar, G. A. C. Murrell, and I. B. Mcinnes, “Inflammatory mechanisms in tendinopathy - towards translation,” Nat. Rev. Rheumatol., vol. 13, no. 2, pp. 110–122, 2017.

[2] M. Franchi, A. Trirè, M. Quaranta, E. Orsini, and V. Ottani, “Collagen structure of tendon relates to function,” ScientificWorldJournal., vol. 7, pp. 404–420, 2007.

[3] M. Benjamin, E. Kaiser, and S. Milz, “Structure-function relationships in tendons: A review,” J. Anat., vol. 212, no. 3, pp. 211–228, 2008.

[4] J. G. Snedeker and J. Foolen, “Tendon injury and repair – A perspective on the basic mechanisms of tendon disease and future clinical therapy,” Acta Biomater., vol. 63, pp. 18–36, 2017.

[5] S. A. Fenwick, B. L. Hazleman, and G. P. Riley, “The vasculature and its role in the damaged and healing tendon,” Arthritis Res., vol. 4, no. 4, pp. 252–260, 2002.

[6] W. Fan, N. Michael, and D. Denitsa, “Tendon injuries: basic science and new repair proposals,” Gen. Orthop., vol. 2, no. July, pp. 211–227, 2017.

[7] D. Docheva, S. A. Müller, M. Majewski, and C. H. Evans, “Biologics for tendon repair,” Adv. Drug Deliv. Rev., vol. 84, pp. 222–239, 2015.

[8] J. Pingel et al., “3-D ultrastructure and collagen composition of healthy and overloaded human tendon: Evidence of tenocyte and matrix buckling,” J. Anat., vol. 224, no. 5, pp. 548–555, 2014.

[9] P. D. Clegg, S. Strassburg, and R. K. Smith, “Cell phenotypic variation in normal and damaged tendons,” Int. J. Exp. Pathol., vol. 88, no. 4, pp. 227–235, 2007.

[10] S. A. Jelinsky, S. A. Rodeo, J. Li, L. V Gulotta, J. M. Archambault, and H. J. Seeherman, “Regulation of gene expression in human tendinopathy,” BMC Musculoskelet Disord, vol. 12, p. 86, 2011.

[11] J. M. Archambault, S. A. Jelinsky, L. S.P., A. Hill, D. Glaser, and L. J. Soslowsky, “Rat Supraspinatus Tendon Expresses Cartilage Markers with Overuse,” J. Orthop. Res., vol. 25, no. May, pp. 1121–1127, 2007.

[12] J. Qi et al., “Differential expression and cellular localization of novel isoforms of the tendon biomarker tenomodulin,” J. Appl. Physiol., vol. 113, no. 6, pp. 861–871, 2012.

[13] N. A. Dyment et al., “The Paratenon Contributes to Scleraxis-Expressing Cells during Patellar Tendon Healing,” PLoS One, vol. 8, no. 3, 2013.

[14] S. Vermeulen et al., “Identification of topographical architectures supporting the phenotype of rat tenocytes,” Acta Biomater., vol. 83, 2019.

[15] L. Yao, C. S. Bestwick, L. A. Bestwick, N. Maffulli, and R. M. Aspden, “Phenotypic drift in human tenocyte culture,” Tissue Eng., vol. 12, no. 7, pp. 1843–1849, 2006.

[16] A. D. Mazzocca et al., “In vitro changes in human tenocyte cultures obtained from proximal biceps tendon: Multiple passages result in changes in routine cell markers,” Knee Surgery, Sport. Traumatol. Arthrosc., vol. 20, no. 9, pp. 1666–1672, 2012.

[17] K. Von Der Mark, V. Gauss, H. Von Der Mark, and P. Müller, “Relationship between cell shape and type of collagen synthesised as chondrocytes lose their cartilage phenotype in culture [26],” Nature, vol. 267, no. 5611, pp. 531–532, 1977.

[18] A. Kraus, D. Sattler, M. Wehland, R. Luetzenberg, N. Abuagela, and M. Infanger, “Vascular endothelial growth factor enhances proliferation of human tenocytes and promotes tenogenic gene expression,” Plast. Reconstr. Surg., vol. 142, no. 5, pp. 1240–1247, 2018.

[19] S. R. Caliari and B. A. C. Harley, “Composite growth factor supplementation strategies to enhance tenocyte bioactivity in aligned collagen-GAG scaffolds,” Tissue Eng. - Part A, vol. 19, no. 9–10, pp. 1100–1112, 2013.

[20] D. Xia et al., “GDFs promote tenogenic characteristics on human periodontal ligament-derived cells in culture at late passages,” Growth Factors, vol. 31, no. 5, pp. 165–173, 2013.

[21] M. Vijven, S. L. Wunderli, K. Ito, J. G. Snedeker, and J. Foolen, “Serum deprivation limits loss and promotes recovery of tenogenic phenotype in tendon cell culture systems,” J. Orthop. Res., no. December 2019, 2020.

[22] S. Webb, C. Gabrelow, J. Pierce, E. Gibb, and J. Elliott, “Retinoic acid receptor signaling preserves tendon stem cell characteristics and prevents spontaneous differentiation in vitro,” Stem Cell Res. Ther., pp. 1–11, 2016.

[23] K. Metavarayuth, P. Sitasuwan, X. Zhao, Y. Lin, and Q. Wang, “Influence of Surface Topographical Cues on the Differentiation of Mesenchymal Stem Cells in Vitro,” ACS Biomater. Sci. Eng., vol. 2, no. 2, pp. 142–151, 2016.

[24] M. Younesi, A. Islam, V. Kishore, J. M. Anderson, and O. Akkus, “Tenogenic Induction of Human MSCs by Anisotropically Aligned Collagen Biotextiles,” Adv. Funct. Mater., pp. 5762–5770, 2014.

[25] R. Peng, X. Yao, and J. Ding, “Effect of cell anisotropy on differentiation of stem cells on micropatterned surfaces through the controlled single cell adhesion,” Biomaterials, vol. 32, no. 32, pp. 8048–8057, 2011.

[26] J. Zhu et al., “The regulation of phenotype of cultured tenocytes by microgrooved surface structure,” Biomaterials, vol. 31, no. 27, pp. 6952–6958, 2010.

[27] V. Kishore, W. Bullock, X. Sun, W. Scott, V. Dyke, and O. Akkus, “Tenogenic differentiation of human MSCs induced by the topography of electrochemically aligned collagen threads,” Biomaterials, vol. 33, no. 7, pp. 2137–2144, 2012.

[28] T. L. Popielarczyk, A. S. Nain, and J. G. Barrett, “Aligned nanofiber topography directs the tenogenic differentiation of mesenchymal stem cells,” Appl. Sci., vol. 7, no. 1, 2017.

[29] T. Tu et al., “Tendon ECM modified bioactive electrospun fibers promote MSC tenogenic differentiation and tendon regeneration,” Appl. Mater. Today, vol. 18, p. 100495, 2020.

[30] S. R. Caliari, D. W. Weisgerber, M. A. Ramirez, D. O. Kelkhoff, and B. A. C. Harley, “The influence of collagen-glycosaminoglycan scaffold relative density and microstructural anisotropy on tenocyte bioactivity and transcriptomic stability,” J. Mech. Behav. Biomed. Mater., vol. 11, pp. 27–40, 2012.

[31] Z. Yin et al., “The regulation of tendon stem cell differentiation by the alignment of nanofibers,” Biomaterials, vol. 31, no. 8, pp. 2163–2175, 2010.

[32] W. Wang et al., “Aligned nanofibers direct human dermal fibroblasts to tenogenic phenotype in vitro and enhance tendon regeneration in vivo,” Nanomedicine, vol. 11, no. 9, pp. 1055–1072, 2016.

[33] M. Younesi, A. Islam, V. Kishore, J. M. Anderson, and O. Akkus, “Tenogenic Induction of Human MSCs by Anisotropically Aligned Collagen Biotextiles,” Adv. Funct. Mater., pp. 5762–5770, 2014.

[34] J. Foolen, S. L. Wunderli, S. Loerakker, and J. G. Snedeker, “Tissue alignment enhances remodeling potential of tendon-derived cells - Lessons from a novel microtissue model of tendon scarring,” Matrix Biol., vol. 65, pp. 14–29, 2018.

[35] R. Schweitzer et al., “Analysis of the tendon cell fate using Scleraxis, a specific marker for tendons and ligaments,” Development, vol. 128, no. 19, pp. 3855–3866, 2001.

[36] H. R. C. Screen, D. E. Berk, K. E. Kadler, F. Ramirez, and M. F. Young, “Tendon functional extracellular matrix,” J. Orthop. Res., vol. 33, no. 6, pp. 793–799, 2015.

[37] Y. J.H. and H. J., “Tendon proteoglycans: Biochemistry and function,” J. Musculoskelet. Neuronal Interact., vol. 5, no. 1, pp. 22–34, 2005.

[38] W. Y. Tong et al., “Functional replication of the tendon tissue microenvironment by a bioimprinted substrate and the support of tenocytic differentiation of mesenchymal stem cells,” Biomaterials, vol. 33, no. 31, pp. 7686–7698, 2012.

[39] N. R. M. Beijer et al., “Dynamic adaptation of mesenchymal stem cell physiology upon exposure to surface micropatterns,” Sci. Rep., vol. 9, no. 1, pp. 1–14, 2019.

[40] G. Schulze-Tanzil et al., “Redifferentiation of dedifferentiated human chondrocytes in high-density cultures,” Cell Tissue Res., vol. 308, no. 3, pp. 371–379, 2002.

[41] M. M. J. Caron et al., “Redifferentiation of dedifferentiated human articular chondrocytes: Comparison of 2D and 3D cultures,” Osteoarthr. Cartil., vol. 20, no. 10, pp. 1170–1178, 2012.

[42] A. J. Mueller, S. R. Tew, O. Vasieva, and P. D. Clegg, “A systems biology approach to defining regulatory mechanisms for cartilage and tendon cell phenotypes,” no. June, pp. 1–14, 2016.

[43] H. Holtzer, J. Abbott, J. Lash, and S. Holtzer, “the Loss of Phenotypic Traits By Differentiated Cells in Vitro, I. Dedifferentiation of Cartilage Cells,” Proc. Natl. Acad. Sci., vol. 46, no. 12, pp. 1533–1542, 1960.

[44] P. R. Tata et al., “Dedifferentiation of committed epithelial cells into stem cells in vivo,” Nature, vol. 503, no. 7475, pp. 218–223, 2013.

[45] R. Pennock et al., “Human cell dedifferentiation in mesenchymal condensates through controlled autophagy,” Sci. Rep., vol. 5, pp. 1–11, 2015.

[46] Q. Fang, D. Chen, Z. Yang, and M. Li, “In vitro and in vivo research on using Antheraea pernyi silk fibroin as tissue engineering tendon scaffolds,” Mater. Sci. Eng. C, vol. 29, no. 5, pp. 1527–1534, 2009.

[47] C. Zhang et al., “Well-aligned chitosan-based ultrafine fibers committed teno-lineage differentiation of human induced pluripotent stem cells for Achilles tendon regeneration,” Biomaterials, vol. 53, pp. 716–730, 2015.

[48] Z. Yin et al., “Electrospun scaffolds for multiple tissues regeneration invivo through topography dependent induction of lineage specific differentiation,” Biomaterials, vol. 44, pp. 173–185, 2015.

[49] E. Chen et al., “An asymmetric chitosan scaffold for tendon tissue engineering: In vitro and in vivo evaluation with rat tendon stem/progenitor cells,” Acta Biomater., vol. 73, pp. 377–387, 2018.

[50] R. D. Cardwell, L. A. Dahlgren, and A. S. Goldstein, “Electrospun fibre diameter, not alignment, affects mesenchymal stem cell differentiation into the tendon/ligament lineage,” J. Tissue Eng. Regen. Med., vol. 8, no. 2014, pp. 937–945, 2012.

[51] Z. Zheng et al., “Alignment of collagen fiber in knitted silk scaffold for functional massive rotator cuff repair,” Acta Biomater., vol. 51, pp. 317–329, 2017.

[52] S. K. Czaplewski, T. L. Tsai, S. E. Duenwald-Kuehl, R. Vanderby, and W. J. Li, “Tenogenic differentiation of human induced pluripotent stem cell-derived mesenchymal stem cells dictated by properties of braided submicron fibrous scaffolds,” Biomaterials, vol. 35, no. 25, pp. 6907–6917, 2014.

[53] K. Zhou et al., “Nanoscaled and microscaled parallel topography promotes tenogenic differentiation of asc and neotendon formation in vitro,” Int. J. Nanomedicine, vol. 13, pp. 3867–3881, 2018.

[54] A. D. Schoenenberger, J. Foolen, P. Moor, U. Silvan, and J. G. Snedeker, “Substrate fiber alignment mediates tendon cell response to inflammatory signaling,” Acta Biomater., vol. 71, pp. 306–317, 2018.

[55] C. Luxenburg and R. Zaidel-Bar, “From cell shape to cell fate via the cytoskeleton — Insights from the epidermis,” Exp. Cell Res., vol. 378, no. 2, pp. 232–237, 2019.

[56] A. English et al., “Substrate topography: A valuable in vitro tool, but a clinical red herring for in vivo tenogenesis,” Acta Biomater., vol. 27, pp. 3–12, 2015.

[57] Y. Sugimoto et al., “Scx+/Scx9+ progenitors contribute to the establishment of the junction between cartilage and tendon/ligament,” Dev., vol. 140, no. 11, pp. 2280–2288, 2012.

[58] M. R. Buckley et al., “Distributions of types I, II and III collagen by region in the human supraspinatus tendon,” Connect. Tissue Res., vol. 54, no. 6, pp. 374–379, 2013.

[59] J. Zhang and J. H. C. Wang, “Mechanobiological response of tendon stem cells: Implications of tendon homeostasis and pathogenesis of tendinopathy,” J. Orthop. Res., vol. 28, no. 5, pp. 639–643, 2010.

[60] G. Nourissat, F. Berenbaum, and D. Duprez, “Tendon injury: From biology to tendon repair,” Nat. Rev. Rheumatol., vol. 11, no. 4, pp. 223–233, 2015.

[61] V. Kishore, W. Bullock, X. Sun, W. S. Van Dyke, and O. Akkus, “Tenogenic differentiation of human MSCs induced by the topography of electrochemically aligned collagen threads,” Biomaterials, vol. 33, no. 7, pp. 2137–2144, 2012.

[62] R. G. Thakar et al., “Cell-shape regulation of smooth muscle cell proliferation,” Biophys. J., vol. 96, no. 8, pp. 3423–3432, 2009.

[63] S. J. Liliensiek, S. Campbell, P. F. Nealey, and C. J. Murphy, “The scale of substratum topographic features modulates proliferation of corneal epithelial cells and corneal fibroblast,” J. Biomed. Mater. Res. Part A, vol. 79, no. 4, pp. 963–73, 2006.

[64] R. J. McMurray et al., “Nanoscale surfaces for the long-term maintenance of mesenchymal stem cell phenotype and multipotency,” Nat. Mater., vol. 10, no. 8, pp. 637–644, 2011.

[65] S. Gronthos, P. J. Simmons, S. E. Graves, and P. G. Robey, “Integrin-mediated interactions between human bone marrow stromal precursor cells and the extracellular matrix,” Bone, vol. 28, no. 2, pp. 174–181, 2001.

[66] R. J. McMurray, M. J. Dalby, and P. M. Tsimbouri, “Using biomaterials to study stem cell mechanotransduction, growth and differentiation,” J. Tissue Eng. Regen. Med., vol. 9, no. 3, pp. 528–539, 2014.

[67] Y. Wang et al., “Integrin subunits alpha5 and alpha6 regulate cell cycle by modulating the chk1 and Rb/E2F pathways to affect breast cancer metastasis,” Mol. Cancer, vol. 10, no. 1, p. 84, 2011.

