## Supplementary file 1 for "Tendon-derived biomimetic surface topographies induce phenotypic maintenance of tenocytes *in vitro*"

**Area:** Number of pixels in the region.

**Compactness:** The mean squared distance of the object's pixels from the centroid divided by the area. A filled circle will have a compactness of 1, with irregular objects or objects with holes having a value greater than 1.

**Solidity:** The proportion of the pixels in the convex hull that are also in the object. Solidity equals 1 is a solid object (no holes) and <1 is an object with holes or irregular boundary

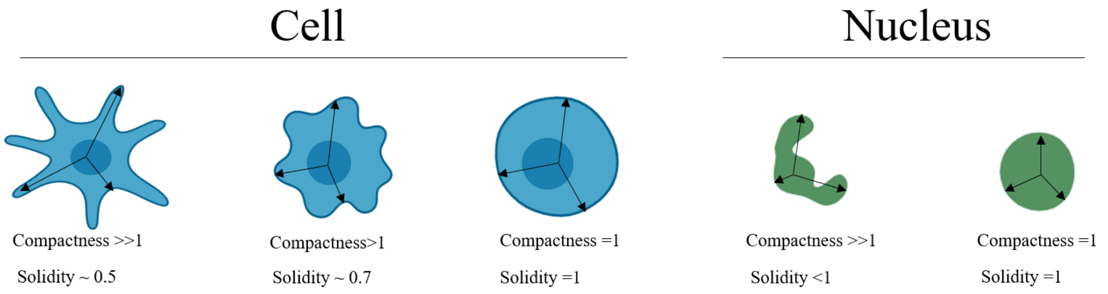


**Eccentricity:** The eccentricity of the ellipse that has the same second-moments as the region. The eccentricity is the ratio of the distance between the foci of the ellipse and its major axis length. The value is between 0 and 1.

**Major/minor axis:** The length of the major axis of the ellipse that has the same normalized second central moments as the region. The length of the minor axis of the ellipse that has the same normalized second central moments as the region. Aspect ratio is major axis divided by minor axis

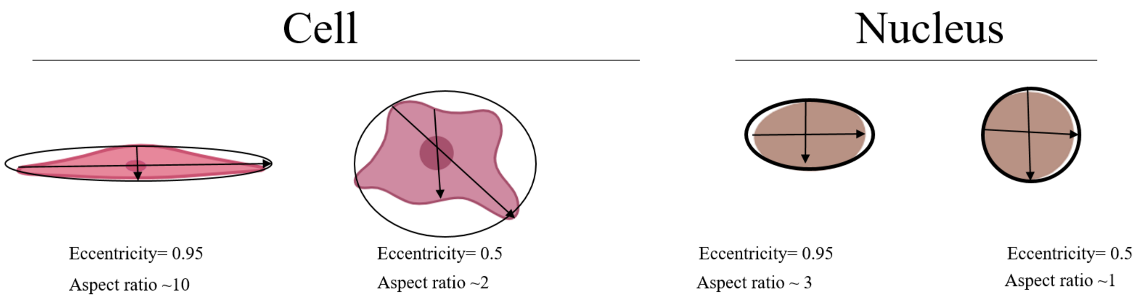


**Supplementary figure 1.** Explanation of shape parameters
